# Proximal tubule cells contribute to the thin descending limb of the loop of Henle during mouse kidney development

**DOI:** 10.1101/2025.01.14.633065

**Authors:** Eunah Chung, Fariba Nosrati, Mike Adam, Andrew Potter, Mohammed Sayed, Christopher Ahn, Benjamin D. Humphreys, Hee-Woong Lim, Yueh-Chiang Hu, S. Steven Potter, Joo-Seop Park

## Abstract

**Background:** The thin descending limb of the loop of Henle is crucial for urine concentration, as it facilitates passive water reabsorption. Despite its importance, little is known about how this nephron segment forms during kidney development.

**Methods:** We assembled a large single-cell RNA sequencing (scRNA-seq) dataset by integrating multiple datasets of non-mutant developing mouse kidneys to identify developing thin descending limb cells. To test whether those cells originate from proximal tubule cells, we generated a proximal tubule-specific Cre line, *Slc34a1eGFPCre*, and conducted lineage tracing. Additionally, given that the transcription factor Hnf4a directly binds to the *Aqp1* gene, we examined whether the loss of Hnf4a affects *Aqp1* expression in thin descending limb cells.

**Results:** From our scRNA-seq dataset, we identified a small cluster of cells distinct from both the proximal tubule and the thick ascending limb of the loop of Henle. Those cells exhibited high expression of thin descending limb marker genes, including *Aqp1* and *Bst1*. Notably, a subset of proximal tubule cells also expressed thin descending limb marker genes, suggesting that proximal tubule cells may give rise to thin descending limb cells. Using lineage tracing with the *Slc34a1eGFPCre* line, we found that, at least, a subset of thin descending limb cells are descendants of proximal tubule cells. Furthermore, the loss of Hnf4a, a transcription factor essential for mature proximal tubule cell formation, disrupted proper *Aqp1* expression in thin descending limb cells, providing additional evidence of a developmental link between proximal tubule cells and thin descending limb cells.

**Conclusion:** Our findings shed new light on the developmental origin of thin descending limb cells and highlight the importance of Hnf4a in regulating their formation.

**Key Points:**

- Reference single cell RNA-seq dataset of the developing mouse kidney was assembled and used to identify the thin descending limb of the loop of Henle.
- Lineage analysis of proximal tubules in the mouse kidney shows that proximal tubule cells give rise to the thin descending limb of the loop of Henle.
- Deletion of Hnf4a disrupts the expression of *Aqp1* in the thin descending limb of the loop of Henle, highlighting a developmental link between proximal tubules and the loop of Henle.

## Introduction

Mesenchymal nephron progenitor cells (mNPs) have the potential to differentiate into any cell type within the nephron.^1^ During development, the mesenchymal-to-epithelial transition of mNPs leads to the formation of the renal vesicle, which develops into the S-shaped body.^2,3^ Although it is known that Wnt,^4–6^ Fgf,^7,8^ and Notch^9,10^ signaling pathways play critical roles in the initial differentiation, it remains poorly understood how the S-shaped body progresses into a fully developed nephron. This is, at least in part, due to the fact that nephron formation is not synchronized in the mammalian kidney. New nephrons form in waves, largely occurring alongside the branching of the ureteric epithelium. This asynchronous development makes it challenging to study the development of the nephron in a stepwise manner.

Single-cell RNA sequencing (scRNA-seq) offers a powerful tool to help overcome this obstacle with its capability to address cellular heterogeneity.^11^ In this study, we leveraged scRNA-seq to investigate how different nephron segments are formed. Our analysis of the mouse kidney reveals that the thin descending limb of the loop of Henle emerges, at least in part, from proximal tubule cells, providing new insights into the developmental origin of the thin descending limb. Additionally, the comprehensive scRNA-seq dataset and the proximal tubule-specific Cre line presented here will serve as valuable resources and tools for kidney research.

## Methods

### Mice

*Slc34a1eGFPCre* was generated by modifying the previously reported *Slc34a1eGFPCreERT2* (JAX: 032285)^12^ by CRISPR.^13^ The guide RNA target sequence (GGCGATCTCGAGCCATCTGC) was selected according to the on- and off-target scores from the web tool CRISPOR (http://crispor.tefor.net).^14^ The chemically modified single-guide RNA (sgRNA) was purchased from IDT. To form ribonucleoprotein complex (RNP), sgRNA was mixed with Cas9 protein (IDT) in Opti-MEM (ThermoFisher) and incubated at 37°C for 10 min. The donor oligo (Ultramer from IDT) with the intended mutations to introduce a stop codon and asymmetrical homologous arm design was added to the RNP.^15^ The final concentrations are 60 ng/μl of sgRNA, 80 ng/μl of Cas9 protein, and 500 ng/μl of donor oligo. The zygotes carrying *Slc34a1eGFPCreERT2* from superovulated female mice were electroporated with 7 μl of the RNP/donor mix on ice using a Genome Editor electroporator (BEX; 30V, 1ms width, and 5 pulses with 1s interval). Two minutes after electroporation, zygotes were moved into 500 μl cold M2 medium (Sigma), warmed up to room temperature, and then transferred into the oviductal ampulla of pseudopregnant CD-1 females. Pups were genotyped by PCR, EcoRI digest, and Sanger sequencing. This strain will be available at The Jackson Laboratory Repository (JAX:040319). Other mouse alleles used in this study have been previously reported: *Rosa26 LacZ* (JAX:003474),^16^ *Hnf4a flox*,^17^ *Osr2Cre* (JAX:009388),^18^ *Rosa26 NuTrap* (JAX:029899),^19^ *Rosa26 Ai3* (JAX: 007903),^20^ and *Six2TGC* (JAX:009606).^1,5^ Animals were housed in a controlled environment with a 12-h light/12-h dark cycle, with free access to water and a standard chow diet. All experiments were performed in accordance with animal care guidelines and the protocols were approved by the Institutional Animal Care and Use Committee of Cincinnati Children’s Hospital Medical Center or Northwestern University. We adhere to the NIH Guide for the Care and Use of Laboratory Animals.

### scRNA-seq

Dissociation was performed using psychrophilic proteases.^21^ Briefly, tissue was digested in the buffer containing 10 mg/ml *Bacillus Licheniformis* enzyme (Sigma, P5380) for 20 min, with trituration and vigorous shaking. The digest mixture was then filtered using a 30 µM filter (Miltenyi). The flow-through was centrifuged at 300g for 5 min at 4°C. Supernatant was removed, and the pellet was resuspended in 1 ml phosphate-buffered saline (PBS) containing 10% FBS, followed by filtration using a 20 µM filter (pluriSelect), and centrifuging the flow-through at 300 g for 5 min at 4°C. The pellet was resuspended in 0.4-1 ml ice-cold 10% FBS/PBS. After the cell suspension was analyzed using a hemocytometer with trypan blue, 9600 cells were loaded into the 10X Chromium instrument for each sample, and Gel Beads in Emulsion (GEMs) were generated. 10X Genomics 3’ v3.1 chemistry was used, using the protocol provided by 10X Genomics, with 14 cycles for cDNA amplification. Single cell libraries were sequenced using the NovaSeq 6000.

### scRNA-seq data analysis

Fastq files were processed through the CellRanger pipeline v6.1.2^22^ using 10X Genomics’ mm10 reference genome, with the setting include introns set to True. Background reads were removed using decontX from the celda package^23^ using the filtered barcodes as the cells to keep and the remaining cells in the raw barcode matrix as the background. Doublets were minimized using DoubletFinder.^24^ The R v4.1.1 library Seurat v4.9^25^ was used for cell type clustering and marker gene identification. Cells expressing >500 genes were retained for downstream analysis. Each sample was normalized SCTransform (v2), using the glmGamPoi method and the number of RNA molecules per cell were regressed out. The effect of cycling cells was regressed out using cell cycle phase scoring. Samples were integrated with the top 3000 common anchor genes to minimize sample to sample variation. Cell clusters were determined by the Louvain algorithm using a resolution of 0.5. UMAP dimension reduction was done using the first thirty principal components. Marker genes for each cell type were calculated using the Wilcoxon Rank Sum test returning only genes that are present in a minimum of 25% of the analyzed cluster. Manual curation of the resulting data was performed by removing cell clusters that were considered junk, clustering only by ribosomal genes, or high mitochondrial gene content. Stromal cells were further subclustered using Seurat’s FindClusters function. To estimate the number of stromal subtypes, clustering was repeated multiple times, each using a different resolution value (ranges from 0.02 to 1, step = 0.02). Then, the resolution value corresponding to the most frequent number of clusters was picked (resolution = 0.36). Marker genes of stromal subtypes were predicted using the same criteria mentioned above with the exception of including only genes whose expression in the analyzed cluster is at least 0.5 log fold-change higher compared to their average expression in other clusters. RNA velocity analysis was performed with epithelial nephron progenitors, early proximal tubules, and proximal tubules. Selected cells were reintegrated using Seurat’s “rpca” method and further doublet removal was done using scDblFinder.^26^ Seurat’s “RunUMAP” was used to run UMAP dimensionality reduction using top 20 principal components. Spliced/unspliced transcript counts were computed from BAM files using velocyto tool.^27^ RNA velocity estimation for these cells was done using scVelo using “stochastic” mode.^28^ Finally, RNA velocity vectors were projected into the UMAP plot.

### Immunofluorescence Staining

Kidneys were fixed in PBS containing 4% paraformaldehyde for 15 min, incubated overnight in 10% sucrose/PBS at 4°C, and embedded in OCT (Fisher Scientific). Cryosections (8-10μm) were incubated overnight with primary antibodies in PBS containing 5% heat-inactivated sheep serum and 0.1% Triton X-100. Primary antibodies used in this study are listed in Supplemental Table 1. Fluorophore-labeled secondary antibodies were used for indirect visualization of the target (see Supplemental Methods). Images were taken with a Nikon Ti-2 inverted widefield microscope equipped with an Orca Fusion camera and Lumencor Spectra III light source.

### Fluorescence activated cell sorting (FACS) and bulk RNA-seq

FACS-isolated cells from *Hnf4a* mutant and control kidneys, where *Osr2Cre* was used to activate *Rosa26 Ai3* reporter and delete *Hnf4a*, were subjected to RNA-seq. Embryonic kidneys at E18.5 were dissociated using TrypLE Select Enzyme 10X (Gibco) and mechanical disruption by repetitive pipetting. Cells were washed and resuspended in PBS containing 1% FBS and 10mM EDTA, then filtered through 40μm Nylon Cell Strainer (BD Falcon). GFP+ cells were isolated using BD FACSAria flow cytometer device. Total RNA from FACS isolated cells were prepared using Single Cell RNA Purification Kit (Norgen) followed by mRNA was isolation using NEBNext Poly(A) mRNA Magnetic Isolation Module (E7490, New England Biolabs). Fragmentation of mRNA and cDNA synthesis was done using NEBNext RNA First Strand Synthesis Module (E7525L) and NEBNext RNA Second Strand Synthesis Module (E6111L) following manufacturer’s instructions. The double-stranded cDNA’s were processed to sequencing libraries using ThruPLEX DNA-seq 12S Kit (R400428, Clontech Laboratories), and the libraries were sequenced on Illumina NextSeq 500.

### Bulk RNA-seq analysis

RNA-seq reads were aligned to UCSC mouse genome mm10 using STAR aligner.^29^ Only uniquely aligned reads were retained for downstream analysis. Raw read counts for each gene were measured using FeatureCounts in the subread package with an option, “-s 2 -O --fracOverlap 0.8”.^30^ Differential gene expression analysis was performed using DESeq2.^31^ Genes with fold-change > 1.5 and FDR < 0.05 were selected as differentially expressed genes.

## Results

Realizing that scRNA-seq datasets with limited cell numbers often fail to show the presence of rare cell types, we assembled a reference scRNA-seq dataset (67,438 cells) by combining seven scRNA-seq datasets of non-mutant mouse kidneys at embryonic day 18.5 (E18.5) or postnatal day 0 (P0) (Figure 1A, Supplemental Figure 1, Supplemental Table 2).^32^ At this stage, active nephrogenesis still occurs while nephron tubule cells with segmental identities are already formed. Therefore, this dataset captures all stages of nephrogenesis, including undifferentiated mNPs, epithelial nephron progenitors (eNPs) present in nascent nephron structures such as renal vesicle and S-shaped body, along with mature nephron segments. Batch correction was successful as the same cell types from each sample clustered together (Supplemental Figure 2).

**Figure 1.**
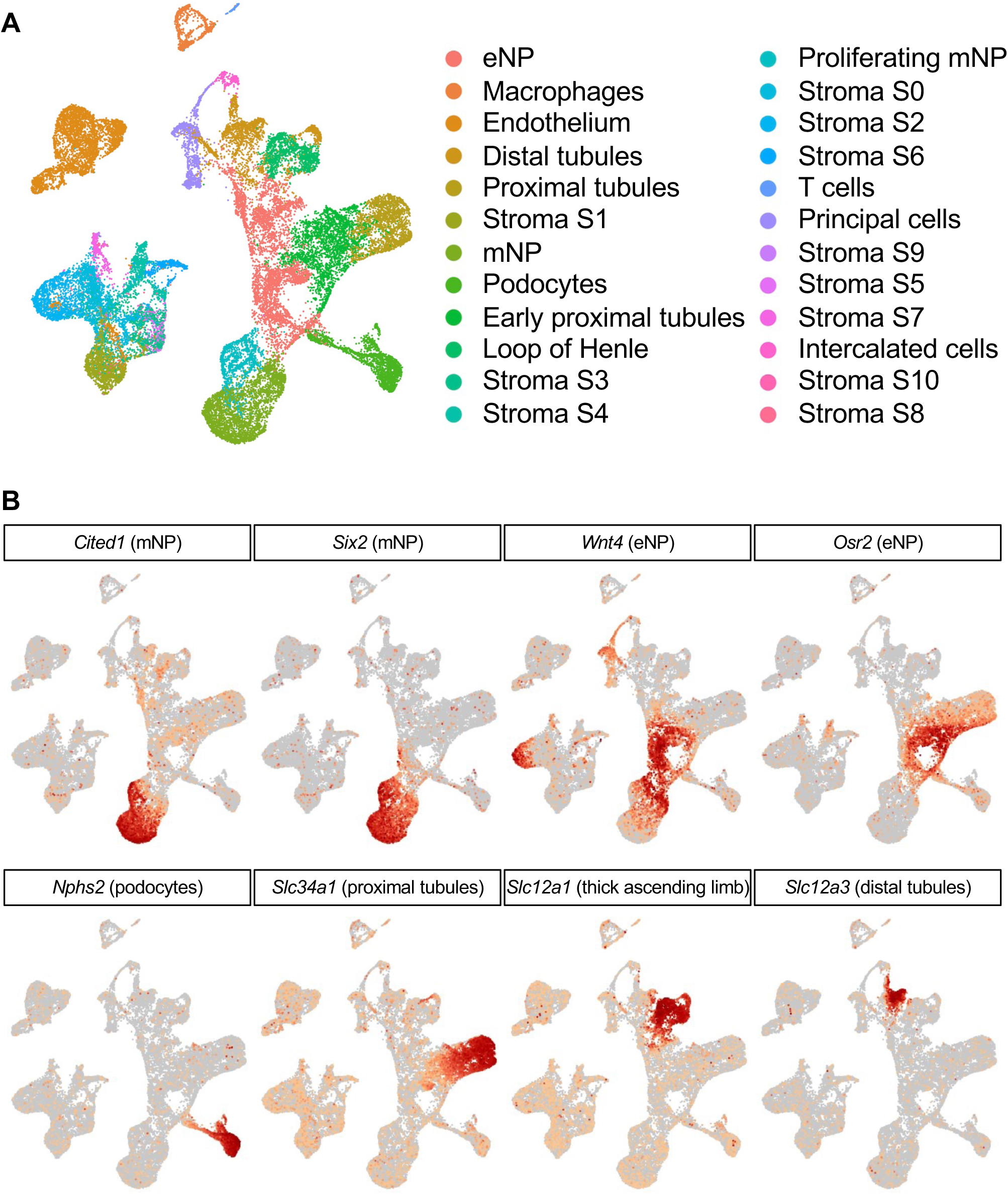
Reference scRNA-seq dataset of the mouse kidney at E18.5 and P0 (A) UMAP plot showing 24 clusters of cells identified in the developing mouse kidney (B) FeaturePlots showing a brief timeline of nephrogenesis. mNPs express *Cited1* and *Six2*, while eNPs express *Wnt4* and *Osr2*. The eNPs develop into four major cell types: podocytes, proximal tubules, thick ascending limb of loop of Henle, and distal tubules.

FeaturePlots in Figure 1B show a brief timeline of mNPs developing into distinct nephron segments. mNPs express *Cited1*^33,34^ and *Six2*,^1,35^ genes that are downregulated when mNPs undergo mesenchymal-to-epithelial transition to become eNPs. During differentiation, *Wnt4*^5,6,36^ and *Osr2*^18^ are sequentially activated. According to previous lineage analyses, *Wnt4Cre* targets all nephron segments^10^ while *Osr2Cre* targets all nephron segments except for the distal tubule.^37^ eNPs eventually develop into the four major cell types of the nephron: podocytes, proximal tubules, loop of Henle, and distal tubules.

Most scRNA-seq studies of mouse kidneys to date have used *Slc12a1* as a marker for loop of Henle,^38,39^ which consists of the thin descending limb, thin ascending limb, and thick ascending limb. However, while *Slc12a1* effectively marks thick ascending limb, it does not capture the entire loop of Henle (Supplemental Figure 3). To identify thin descending limb cells in our scRNA-seq dataset, we analyzed genes specifically expressed in the thin descending limb using previously reported bulk RNA-seq data generated from microdissected nephron segments of adult mouse kidneys (GSE150338).^40^ Consistent with a previous report,^41^ we found that *Bst1* and *Aqp1* were preferentially expressed in the thin descending limb (Supplemental Table 3). FeaturePlots revealed that both genes were strongly expressed in a small cluster adjacent to the proximal tubule cluster (black arrows in Figure 2A). Notably, this small cluster with high *Bst1* and *Aqp1* expression showed little or no expression of the proximal tubule marker *Slc34a1* or the thick ascending limb marker *Slc12a1* (Figure 1B, for magnified view see Supplemental Figure 4), distinguishing it from proximal tubule or thick ascending limb cells. These observations suggest that this cluster likely represents thin descending limb cells.

**Figure 2.**
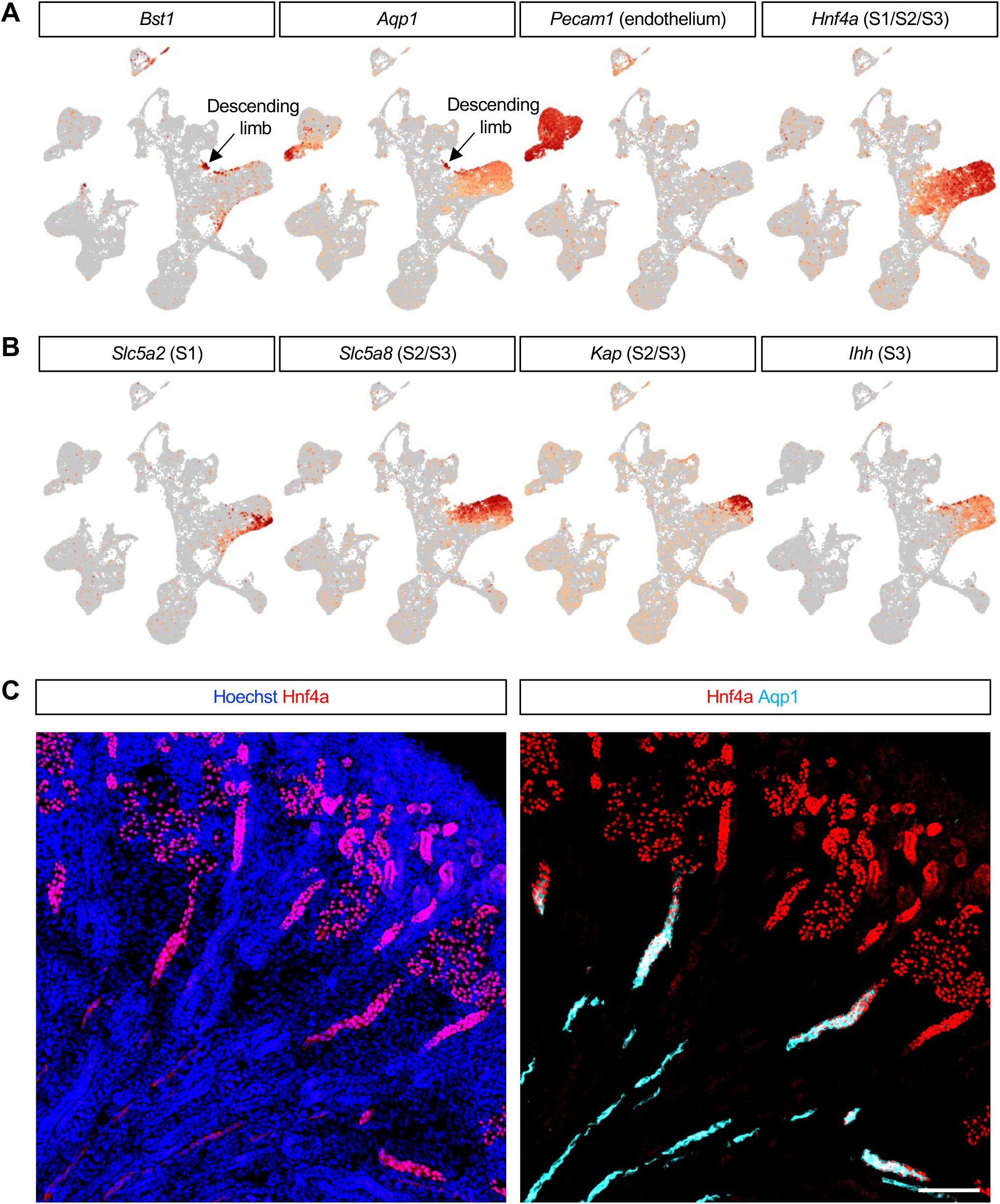
Identification of thin descending limb cells from scRNA-seq dataset (A) FeaturePlots showing the expression of thin descending limb marker genes *Bst1* and *Aqp1*. Black arrow marks the cluster representing thin descending limb cells. *Aqp1* is expressed not only in thin descending limb cells but also in a subset of endothelial cells. FeaturePlots for *Pecam1* and *Hnf4a* mark the clusters for endothelium and proximal tubule cells, respectively. (B) FeaturePlots showing subsegments of proximal tubule cells. *Slc5a2* and *Ihh* mark S1 and S3 segments, respectively. While both *Slc5a8* and *Kap* mark S2/S3 segments, the activation of *Slc5a8* appears to proceed prior to that of *Kap*. (C) Aqp1 marks Hnf4a+ straight tubules but it is undetectable in convoluted proximal tubule cells in the mouse embryonic kidney. Stage, E18.5; Scale Bar, 100μm.

Interestingly, both *Bst1* and *Aqp1* were also detectable at the upper edge of the proximal tubule cluster (Figure 2A, Supplemental Figure 4). The proximal tubule segment can be divided into three subsegments: the convoluted S1 and S2 segments and the straight S3 segment. In the developing mouse kidney, the S1 segment appeared distinct from the S2/S3 segments (Figure 2B). Genes specific to S1, such as *Slc5a2*, were expressed in the lower part of the proximal tubule cluster, while S2/S3-specific genes, such as *Slc5a8* and *Kap*, were expressed in the upper part.^40^ Notably, *Ihh*, which has been shown to be expressed in the straight proximal tubule tubules (S3 segment) of the developing kidney,^42^ exhibited strong expression at the upper edge of the proximal tubule cluster (Figure 2B, Supplemental Figure 4). This expression pattern was intriguing, as both thin descending limb marker genes, *Bst1* and *Aqp1*, appeared to be expressed in the S3 segment of proximal tubule (Figure 2A, Supplemental Figure 4). To validate this finding, we performed immunostaining. As shown in Figure 2C, Hnf4a*+* straight tubules (S3 segment) of the developing mouse kidney were also positive for Aqp1. Since *Aqp1* is continuously expressed in thin descending limb cells, this result raises the intriguing possibility that proximal tubule cells in the S3 segment may give rise to thin descending limb cells.

In order to test if thin descending limb cells originate from proximal tubule cells, we generated a new proximal tubule-specific Cre line. The expression of *Slc34a1* is highly specific for proximal tubule cells (Figure 1B, Supplemental Figure 4).^40^ We converted the tamoxifen-inducible *Slc34a1eGFPCreERT2*^12^ to a constitutively active Cre (*Slc34a1eGFPCre*) by inserting a stop codon immediately after the Cre coding sequence (Supplemental Figure 5). With this new Cre line, we performed lineage tracing of proximal tubule cells. In the adult mouse kidney, convoluted proximal tubule cells (S1 and S2 segments) exhibited strong LTL staining, whereas straight proximal tubule tubules (S3 segment) showed weaker LTL staining (Figure 3A). All proximal tubule cells across these segments expressed *Hnf4a*. When the *Rosa26 Sun1* reporter was activated by *Slc34a1eGFPCre*, the reporter was expressed in most Hnf4a*+* cells in the S1 and S2 segments, as well as in a subset of Hnf4a*+* cells in the S3 segment (Figure 3A). This indicates that *Slc34a1Cre* targets all three proximal tubule segments with minor mosaicism in the S3 segment. Additionally, the reporter was activated in Hnf4a*-* cells located in the medulla. These cells were identified as thin descending limb cells based on their positive expression of *Aqp1* and lack of *Slc12a1* (Figure 3B). This result indicates that thin descending limb cells are descendants of proximal tubule cells. This finding was also supported by our RNA velocity analysis suggesting that Aqp1+ descending limb emerge, at least in part, from Slc34a1+ proximal tubules (Supplemental Figure 7).

**Figure 3.**
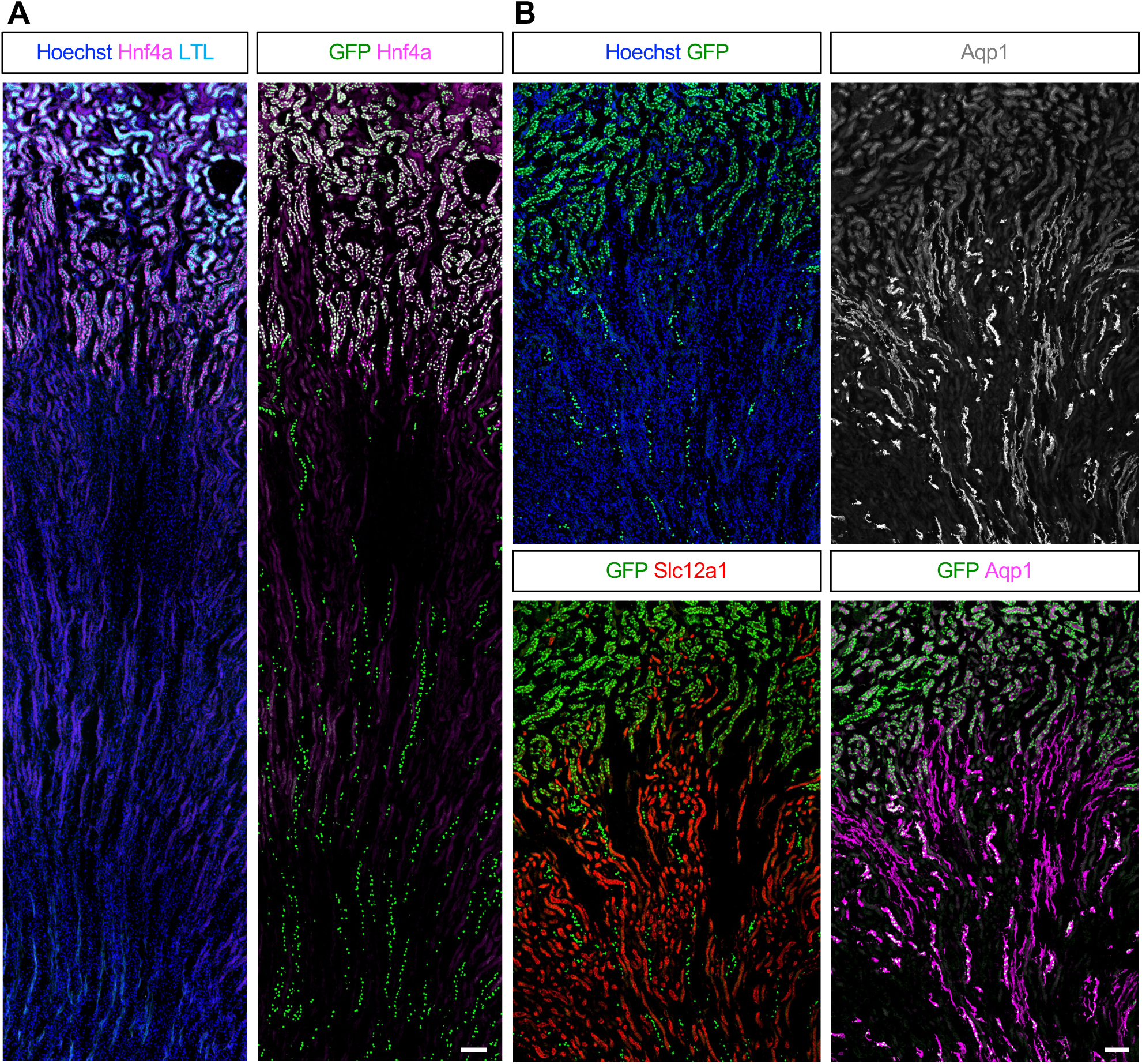
Lineage tracing of proximal tubule cells using *Slc34a1eGFPCre* shows that proximal tubule cells give rise to Aqp1+ thin descending limb cells (A) Cre-dependent activation of *Rosa26 Sun1* reporter is detectable not only in Hnf4a+ proximal tubule cells at the cortex but also in Hnf4a- cells at the medulla. (B) In the adult mouse kidney, Aqp1 is present in both proximal tubule cells and thin descending limb cells although thin descending limb cells exhibit stronger Aqp1 stain than proximal tubule cells. The cells with active *Rosa26 Sun1* reporter in the medulla are positive for Aqp1 but negative for Slc12a1, suggesting that those cells are indeed thin descending limb and not thick ascending limb. Stage, 4 month; Scale Bar, 100μm.

The lineage tracing analysis with *Slc34a1eGFPCre* in newborn mouse kidneys showed that this Cre does not target non-proximal tubule segments such as podocytes or distal tubules (Supplemental Figure 6A). Furthermore, *Slc34a1eGFPCre*-mediated activation of *Rosa26 Sun1* reporter recapitulated the endogenous expression of *Slc34a1*, suggesting that *Slc34a1eGFPCre* is specific for proximal tubules (Supplemental Figure 6B). Finally, Sun1+ descending limb cells were negative for Slc34a1, which confirmed that they originated from proximal tubules (Supplemental Figure 6B). These findings suggest that the low expression of *Slc34a1* in the cluster representing the thin descending limb (Supplemental Figure 4) likely results from noisy gene expression, potentially due to incomplete removal of doublets or the presence of free-floating RNA contaminating single-cell libraries.

Our finding that proximal tubule cells give rise to thin descending limb cells led us to investigate the potential role of Hnf4a in thin descending limb formation. Previously, we showed that Hnf4a is required for the formation of mature proximal tubule cells.^43,44^ Moreover, our chromatin immunoprecipitation analysis of Hnf4a in the developing mouse kidneys showed that Hnf4a directly binds to the promoter of *Aqp1*, a gene highly expressed in thin descending limb (Supplemental Figure 8A).^44^ In the mouse kidney lacking Hnf4a, *Aqp1* expression was reduced by 50% (GSE144772),^44^ with the remaining *Aqp1* expression in the *Hnf4a* mutant kidneys likely coming from endothelial cells (Figure 2A). These observations suggest that Hnf4a may play a regulatory role in *Aqp1* expression in the thin descending limb.

Nephron segments are spatially organized along the cortico-medullary axis of the kidney. The loop of Henle is located in the medulla and papilla of the kidney while the other nephron segments are strictly located at the cortex. This is achieved, at least in part, by elongation of loop of Henle through the boundary of the cortex and medulla (Supplemental Figure 3).^45^ To examine loop of Henle formation in the *Hnf4a* mutant and control kidneys, we examined the cells in the nephron lineage located in the medulla of the kidney by performing lineage tracing with *Osr2Cre*, which targets all nephron segments except for the distal tubule.^37^ We found that both control and mutant kidneys formed nephron tubules in the medulla of the kidney and that both kidneys formed Slc12a1+ thick ascending limb (Figure 4A). This result suggests that the absence of Hnf4a does not affect the elongation of the loop of Henle nor the formation of the thick ascending limb.

**Figure 4.**
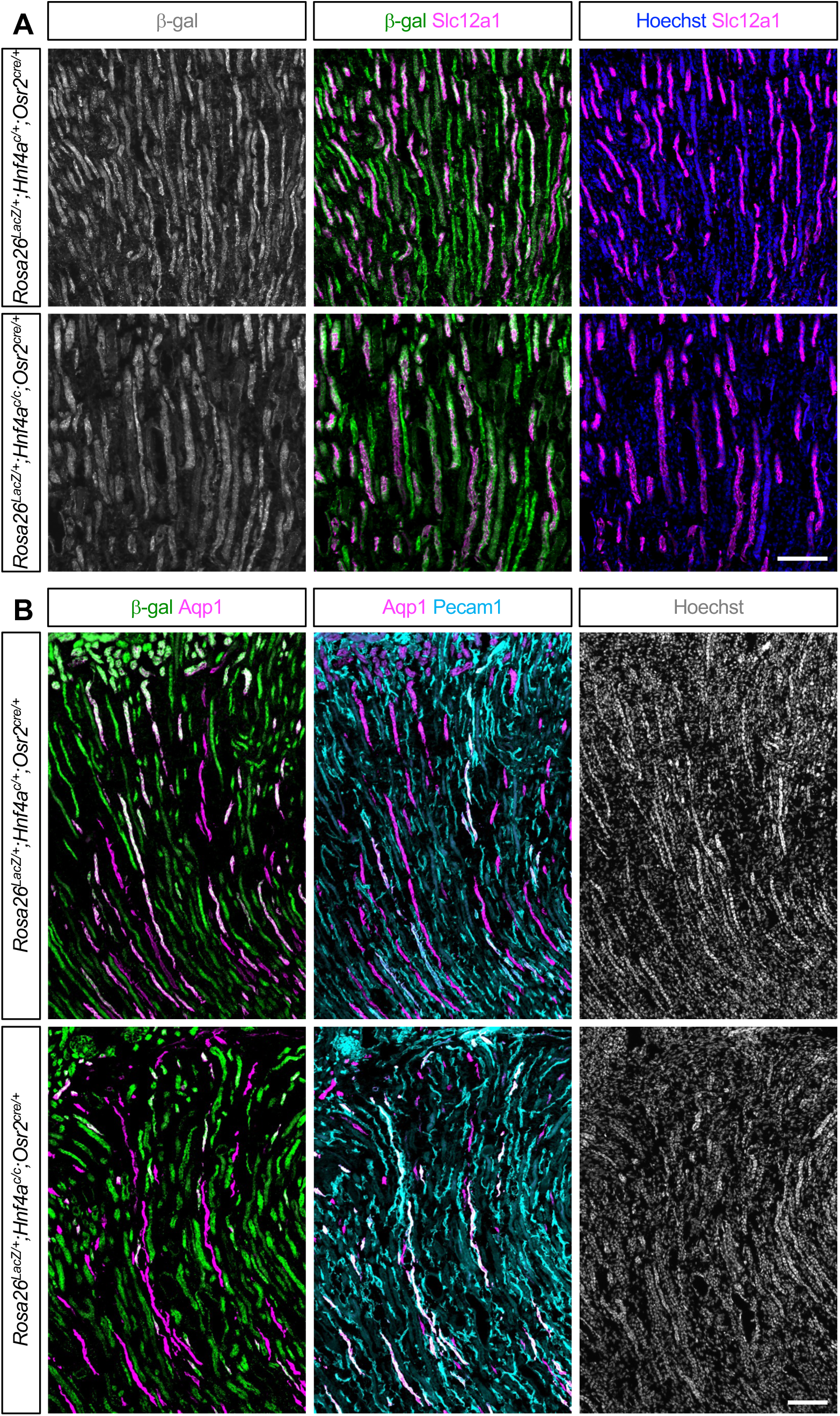
Deletion of *Hnf4a* disrupts *Aqp1* expression in thin descending limb cells. (A) *Osr2Cre*-mediated activation of *Rosa26 LacZ* reporter marks all loop of Henle cells in the medulla. Both control and mutant kidneys have β-galactosidase+ straight tubules and a subset of those β-galactosidase+ cells are also positive for Slc12a1, suggesting that the absence of Hnf4a does not block the elongation of the loop of Henle or the formation of Slc12a1+ thick ascending limb cells. (B) Given *Aqp1* is expressed in both thin descending limb and a subset of the endothelium (Figure 2A), Aqp1+ cells in the control kidney are positive for either β- galactosidase or Pecam1. In the *Hnf4a* mutant kidney, most of the Aqp1+ tubules are positive for Pecam1, suggesting that they are all endothelial cells. A few clusters of β-galactosidase+ Aqp1+ tubules are present but they remain short, suggesting that Hnf4a is required for the proper expression of *Aqp1* in thin descending limb cells. Stage, P10; Scale Bar, 100μm.

In the developing mouse kidney, *Aqp1* is expressed in both thin descending limb cells and a subset of endothelial cells (Figure 2A). To distinguish between these two *Aqp1+* cell populations, we performed co-staining for *Aqp1* along with the *Rosa26 LacZ* lineage tracer activated by *Osr2Cre* and the endothelial marker *Pecam1* (Figure 4B). In the control kidney, all *Aqp1+* straight tubules overlapped with either β-galactosidase or *Pecam1*. Specifically, β-galactosidase+ *Aqp1+* cells represented thin descending limb cells, while *Pecam1+ Aqp1+* cells represented endothelial cells. In the *Hnf4a* mutant kidney, however, most of the *Aqp1+* long tubules were also positive for *Pecam1*, indicating endothelial cells. While β-galactosidase+ *Aqp1+* cells were still present in the mutant kidney, *Aqp1* expression was restricted to a short segment of tubules (Figure 4B). Quantification of the images showed that Aqp1 signal in the thin descending limb in the *Hnf4a* mutant kidney was reduced by 73% (Supplemental Figure 8B). Furthermore, we performed nephron lineage-specific bulk RNA-seq analysis with FACS-isolated cells from the *Hnf4a* mutant and control kidney by *Osr2Cre* at E18.5 (Supplemental Table 4). We found that the loss of Hnf4a reduced *Aqp1* expression by 93%, with no effect on the expression of Slc12a1 (Supplemental Figure 8C), suggesting that Hnf4a is critical for *Aqp1* expression in proximal tubules and thin descending limb. In summary, we found that thin descending limb cells in the *Hnf4a* mutant kidney show defective expression of *Aqp1*, highlighting the critical role of Hnf4a in nephron segmentation beyond proximal tubule cells.

## Discussion

In this study, we provide new insights into the developmental origin of the thin descending limb in the loop of Henle. By analyzing a large integrated scRNA-seq dataset, we identified a distinct cluster of cells with high expression of thin descending limb marker genes, namely *Aqp1* and *Bst1*. Our lineage tracing experiments using the proximal tubule-specific *Slc34a1eGFPCre* line further confirmed that, at least, a subset of thin descending limb cells are descendants of proximal tubule cells. Additionally, we demonstrated that the transcription factor Hnf4a, which plays a critical role in proximal tubule maturation,^43,44^ is also essential for proper *Aqp1* expression in thin descending limb cells. These findings collectively highlight a developmental connection between proximal tubule and thin descending limb cells and reveal an important regulatory role for Hnf4a in thin descending limb cell differentiation.

The assembly of a large scRNA-seq dataset allowed us to identify thin descending limb cells, a rare cell type whose presence has been elusive in previous single-cell studies of the developing mouse kidney. It is unclear why the presence of thin descending limb cells is harder to detect in scRNA-seq data than that of thick ascending limb cells. One possible explanation is that the dissociation procedure used to generate single-cell suspensions introduces biases in cellular composition. Some cell types, such as thin descending limb cells, may be more sensitive to enzymatic or mechanical dissociation, resulting in loss or damage during sample preparation. As a result, the cell diversity observed in scRNA-seq data may not accurately represent the true *in vivo* cellular composition.

Aqp1 and Slc12a1 not only serve as markers for thin descending limb and thick ascending limb, respectively, but also play critical physiological roles in the loop of Henle.^46,47^ Slc12a1 (also known as NKCC2) in the thick ascending limb actively reabsorbs sodium, potassium, and chloride ions from the filtrate traveling the lumen of the nephron, contributing to the hyperosmotic gradient in the medulla of the kidney.^48^ Aqp1 in the thin descending limb allows passive water reabsorption driven by the osmotic gradient established by the thick ascending limb.^49^ This coordinated expression of transporters and channels along the loop of Henle segments establishes and maintains the medullary osmotic gradient, enabling the kidney to concentrate urine via a mechanism known as countercurrent multiplication.^50,51^

The observation that *Aqp1*, a thin descending limb marker gene, is also expressed in the S3 segment of proximal tubule cells suggested a developmental connection between proximal tubule and thin descending limb cells. Our lineage tracing analysis using the proximal tubule-specific *Slc34a1eGFPCre* provided strong evidence that proximal tubule cells contribute to the thin descending limb. Consistent with this, we found that Hnf4a plays a critical role in regulating *Aqp1* expression in the thin descending limb cells, despite the fact that thin descending limb cells exhibit little to no expression of *Hnf4a*. Interestingly, while *Aqp1* is weakly expressed in convoluted proximal tubule cells of the S1/S2 segments, the more robust *Aqp1* expression seen in thin descending limb cells appears to require more than the direct binding of Hnf4a to the *Aqp1* gene. This raises the possibility that Hnf4a may function as a pioneering factor, as observed in other cell types,^52,53^ by making the *Aqp1* gene accessible to another, yet unidentified, regulatory factor.

Bulk RNA-seq analysis of microdissected mouse tubule segments showed that *Slc34a1* expression is specific to the S1 and S2 segments of the proximal tubules in adult mouse kidneys (Supplemental Figure 7E).^40^ Consistent with this, *Slc34a1eGFPCreERT2* primarily targets the S1 and S2 segments but targets the S3 segment poorly even after multiple tamoxifen injections.^12^ This suggests that *Slc34a1* expression is specific to the convoluted proximal tubules. Unlike the inducible *Slc34a1eGFPCreERT2* that is typically used to target proximal tubules in the adult kidney, our constitutively active *Slc34a1eGFPCre* targets all Slc34a1+ cells and their descendants as soon as Slc34a1+ cells appear in the developing kidney. We found that *Slc34a1eGFPCre* targets not only the S1 and S2 segments of the proximal tubules but also the S3 segment and the thin descending limb of the loop of Henle. This result suggests that in the developing mouse kidney, at least, a subset of the S3 segment and descending limb originate from Slc34a1+ proximal tubules. Given that the lineage trace with *Slc34a1GFPCreERT*2 in the adult kidney is restricted to the convoluted proximal tubules, the difference we see with the constitutive *Slc34a1eGFPCre* suggests that the immature proximal tubules during development have the multipotency to develop into the descending limb. It is noteworthy that the targeting of the S3 segment by *Slc34a1eGFPCre* is mosaic, particularly the very distal part of the S3 segment that borders the descending limb (Figure 3A). Having a lineage tracer to distinguish Cre-targeted cells from non-targeted cells would be important. Mosaicism of the descending limb by *Slc34a1eGFPCre* may be due to mosaic targeting of the S3 segment, a putative progenitor population for the descending limb. Alternatively, as our RNA velocity analysis suggests, in addition to the Slc34a1+ cells, another progenitor population (Wnt7b+ Scel+ eNPs) may also contribute to the descending limb (Supplemental Figure 7). Further investigation is necessary to differentiate these possibilities.

In summary, our study establishes a previously unrecognized developmental link between proximal tubule cells and thin descending limb cells, advancing our understanding of nephron segmentation during kidney development. While our findings provide important insights, further studies are needed to uncover the molecular mechanisms driving the transition from the proximal tubule to the thin descending limb with the downregulation of Hnf4a.

## Supporting information

SuppTable2

SuppTable3

SuppTable1

SuppTable4

## Author Contributions

Conceptualization: Eunah Chung, Joo-Seop Park

Data curation: Mike Adam, Joo-Seop Park

Formal analysis: Mike Adam, Mohammed Sayed, Christopher Ahn, Hee-Woong Lim

Funding acquisition: S. Steve Potter, Joo-Seop Park

Investigation: Eunah Chung, Fariba Nosrati, Joo-Seop Park

Methodology: Eunah Chung, Fariba Nosrati, Andrew Potter, Benjamin D. Humphreys, Yueh-Chiang Hu

Project administration: Joo-Seop Park

Resources: Benjamin D. Humphreys, S. Steve Potter, Joo-Seop Park

Software: Mike Adam, Mohammed Sayed, Christopher Ahn, Hee-Woong Lim

Validation: Eunah Chung, Fariba Nosrati, Joo-Seop Park

Visualization: Eunah Chung, Mike Adam, Joo-Seop Park

Writing - original draft: Eunah Chung, Joo-Seop Park

Writing - review & editing: Eunah Chung, Mike Adam, Joo-Seop Park

## Data Sharing Statement

The scRNA-seq and bulk RNA-seq data generated in this study were deposited at the Gene Expression Omnibus (GEO) under accession numbers GSE202882, GSE275601, and GSE291740.

## Supplemental Information

**Supplemental Figure 1.**
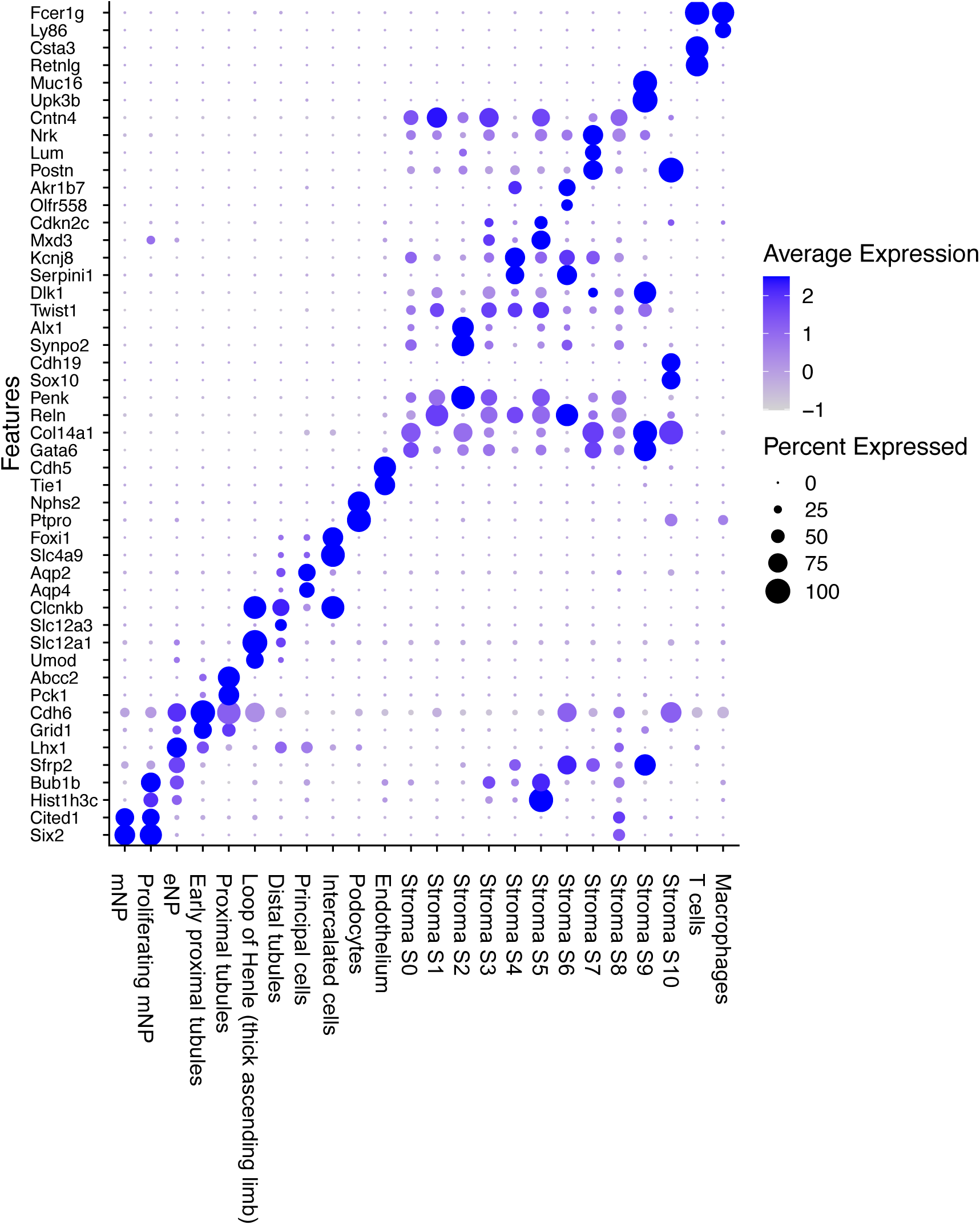
Dot plots showing the expression of representative markers for the clusters identified in scRNA-seq dataset. The full list of the marker genes is presented in Supplemental Table 2.

**Supplemental Figure 2.**
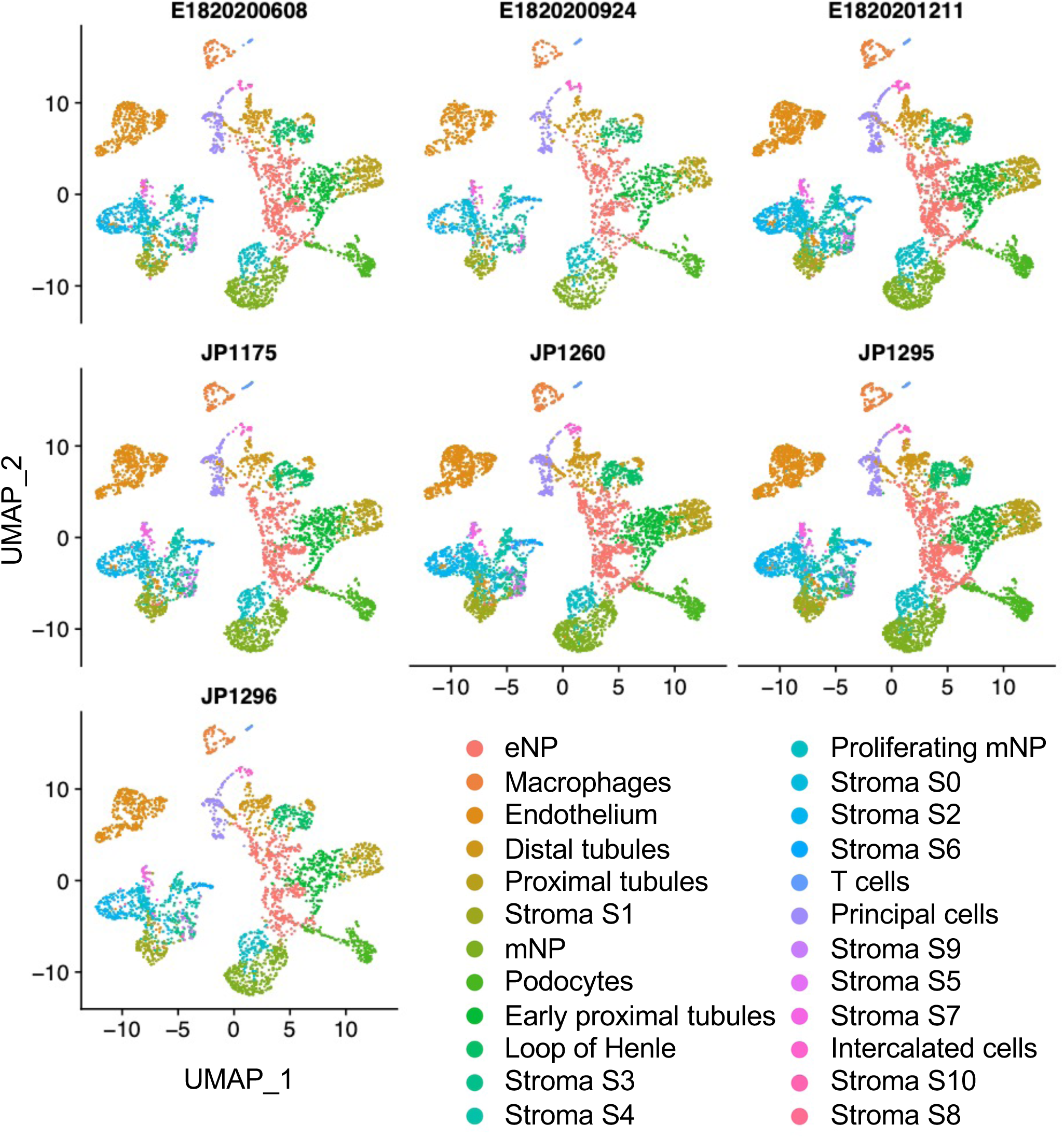
UMAP split by sample. UMAP embedding of scRNA-seq data from seven samples of E18 and P0 mouse kidneys. Each figure shows the cell type distribution of each sample after batch correction. All cell types are present in every sample with none of the samples showing an over or underrepresentation of any cell type.

**Supplemental Figure 3.**
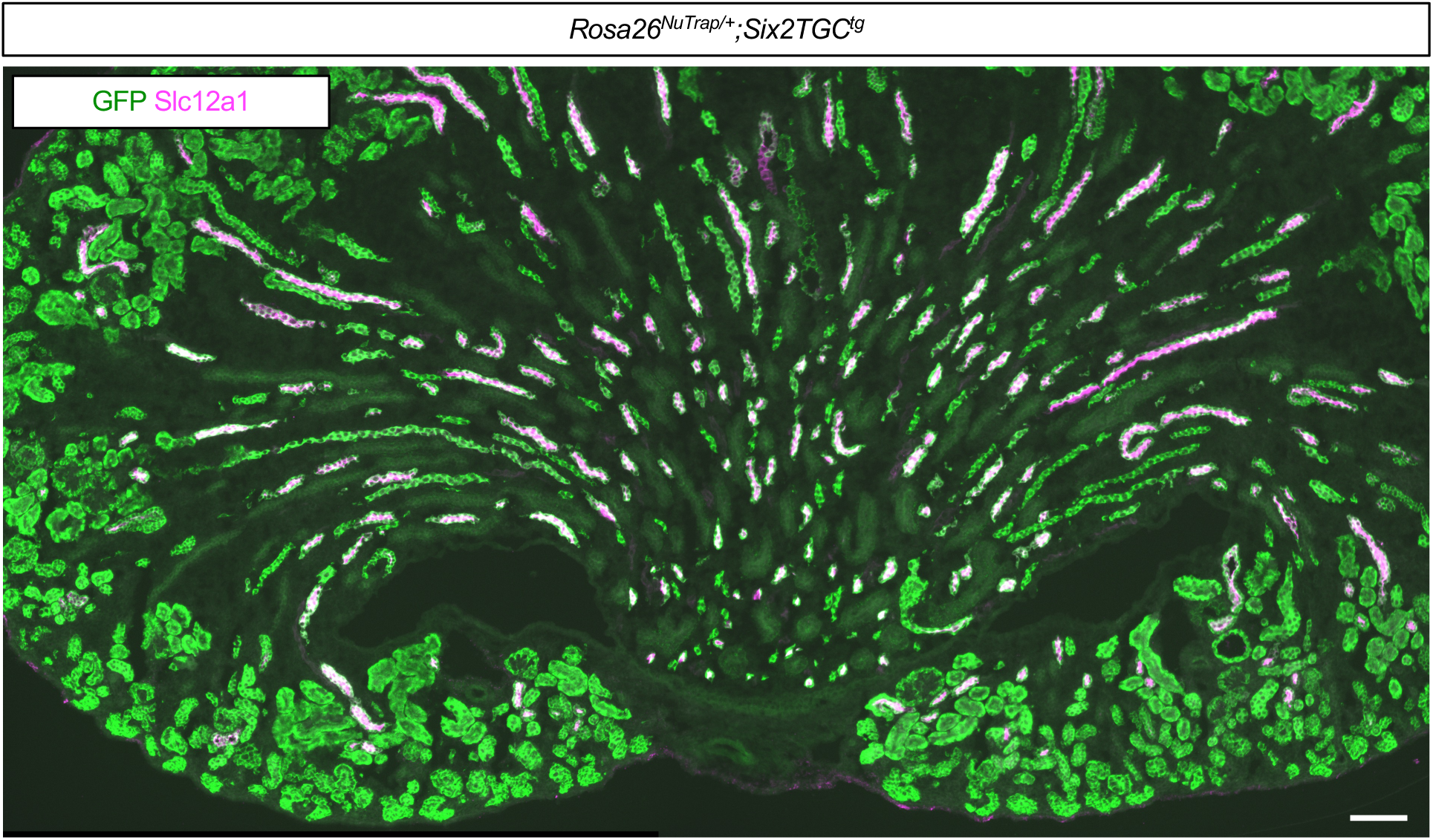
Slc12a1 does not mark all cells in the loop of Henle. GFP reporter (Rosa26-NuTrap) was activated by nephron lineage-specific *Six2TGC*. GFP+ cells in the medulla and papilla represent the loop of Henle. Stage, P0; Scale Bar, 100μm.

**Supplemental Figure 4.**
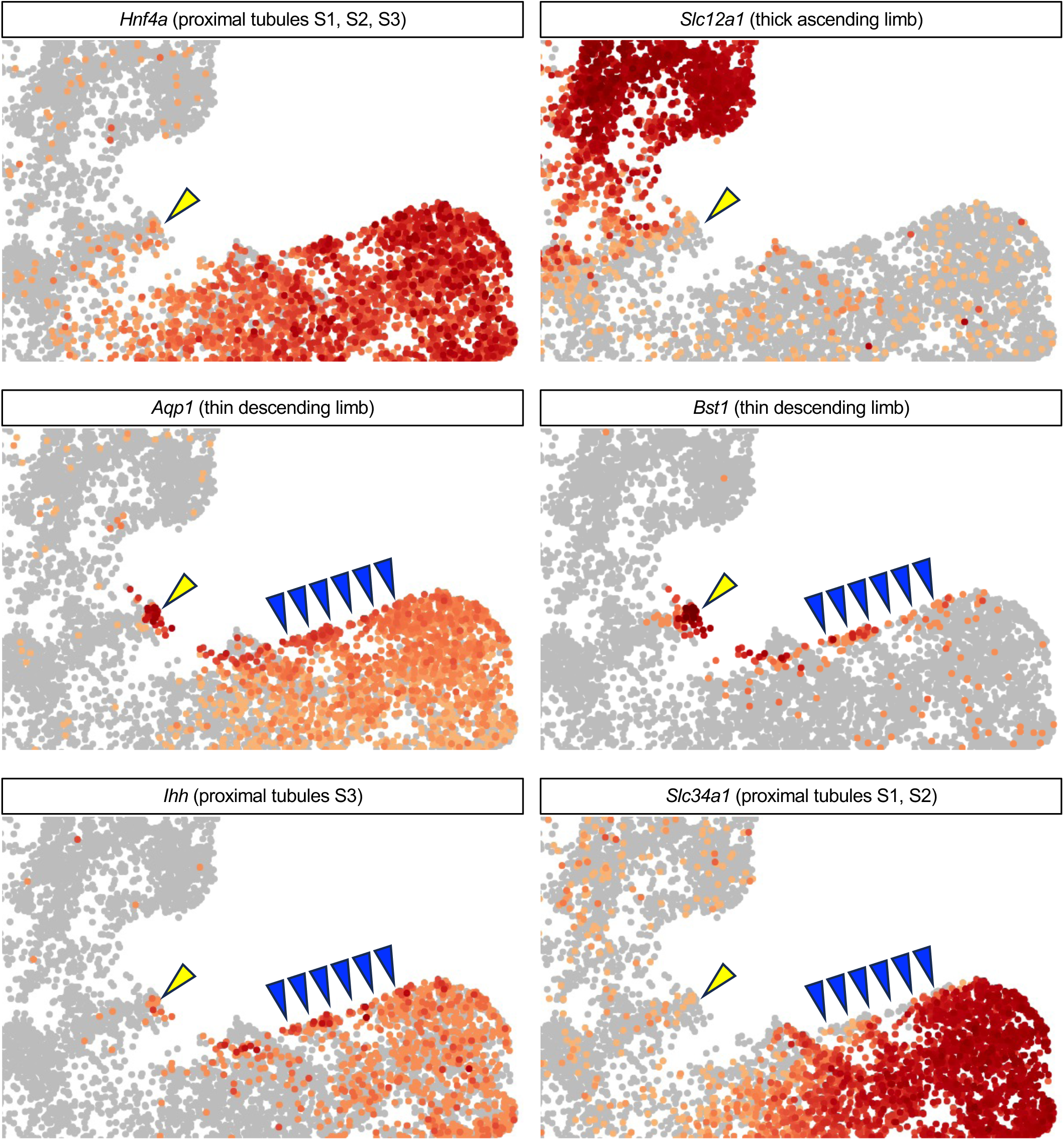
Magnified FeaturePlots show *Hnf4a* is expressed in all three segments (S1, S2, and S3) of proximal tubules while *Slc12a1* is expressed in the thick ascending limb of loop of Henle. Their expression is low or absent in the thin descending limb of loop of Henle (yellow arrowhead). *Aqp1* and *Bst1* are highly expressed in the thin descending limb of loop of Henle (yellow arrowhead) and moderately expressed in the S3 segment of proximal tubules (the upper outline of the proximal tubule cluster, marked by blue arrowheads). *Ihh* is highly expressed in the S3 segment while *Slc34a1* is largely absent in the S3 segment and the thin descending limb.

**Supplemental Figure 5.**
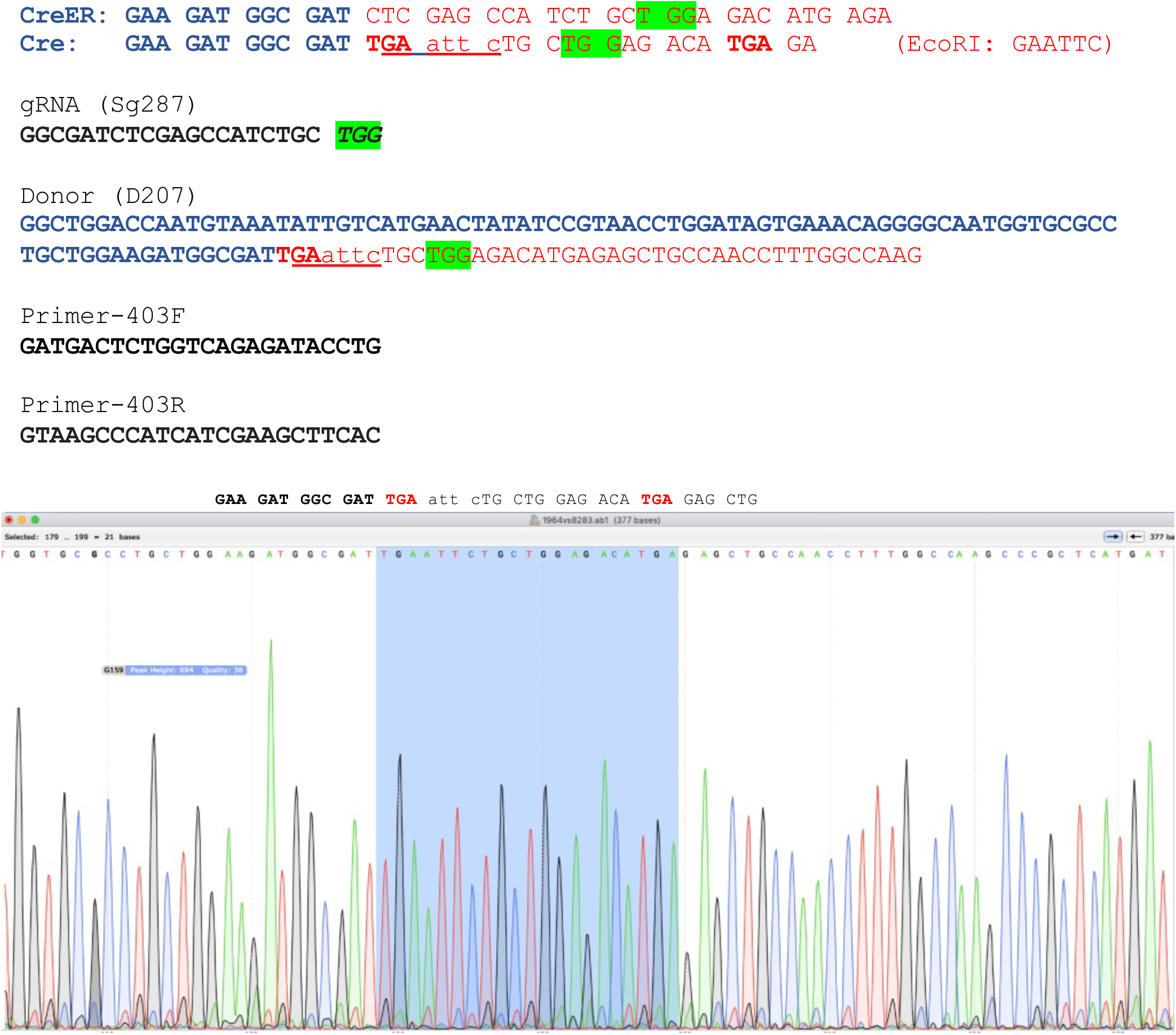
Conversion of a tamoxifen-inducible CreERT2 to a constitutively active Cre using CRISPR. A stop codon (TGA) and an EcoRI (GAATTC) site was added after the coding sequence of Cre recombinase. Green highlight indicates PAM (protospacer adjacent motif) sequence. The 403 bp region was amplified with two primers and diagnostic digestion with EcoRI was performed to identify the successful integration of the doner DNA. The correct targeting was confirmed by Sanger sequencing.

**Supplemental Figure 6.**
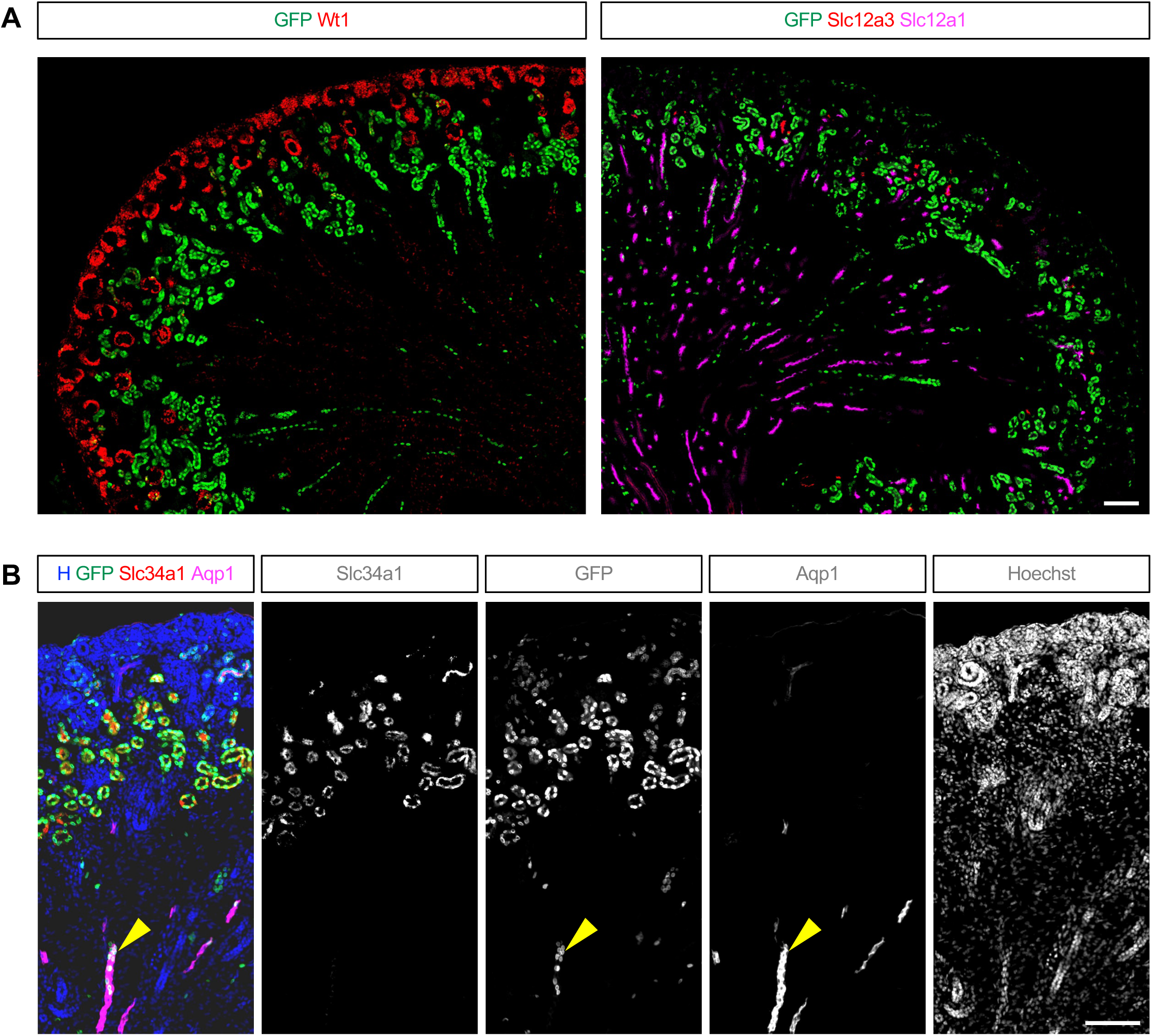
The specificity of *Slc34a1eGFPCre* at P1 mouse kidney. **(A)** *Slc34a1eGFPCre*-mediated activation of *Rosa26 Sun1* reporter shows that *Slc34a1eGFPCre* largely do not target Wt1+ nephron progenitors and podocytes, Slc12a1+ thick ascending limb of the loop of Henle, and Slc12a3+ distal tubules. **(B)** *Slc34a1eGFPCre*-mediated activation of *Rosa26 Sun1* reporter shows good correlation with the endogenous expression of *Slc34a1* in the cortex. A subset of Aqp1+ cells (yellow arrowhead) are positive for Sun1 reporter and negative for Slc34a1, suggesting that they emerge from Slc34a1+ proximal tubules located in the cortex. Stage, P1; Scale Bar, 100μm.

**Supplemental Figure 7.**
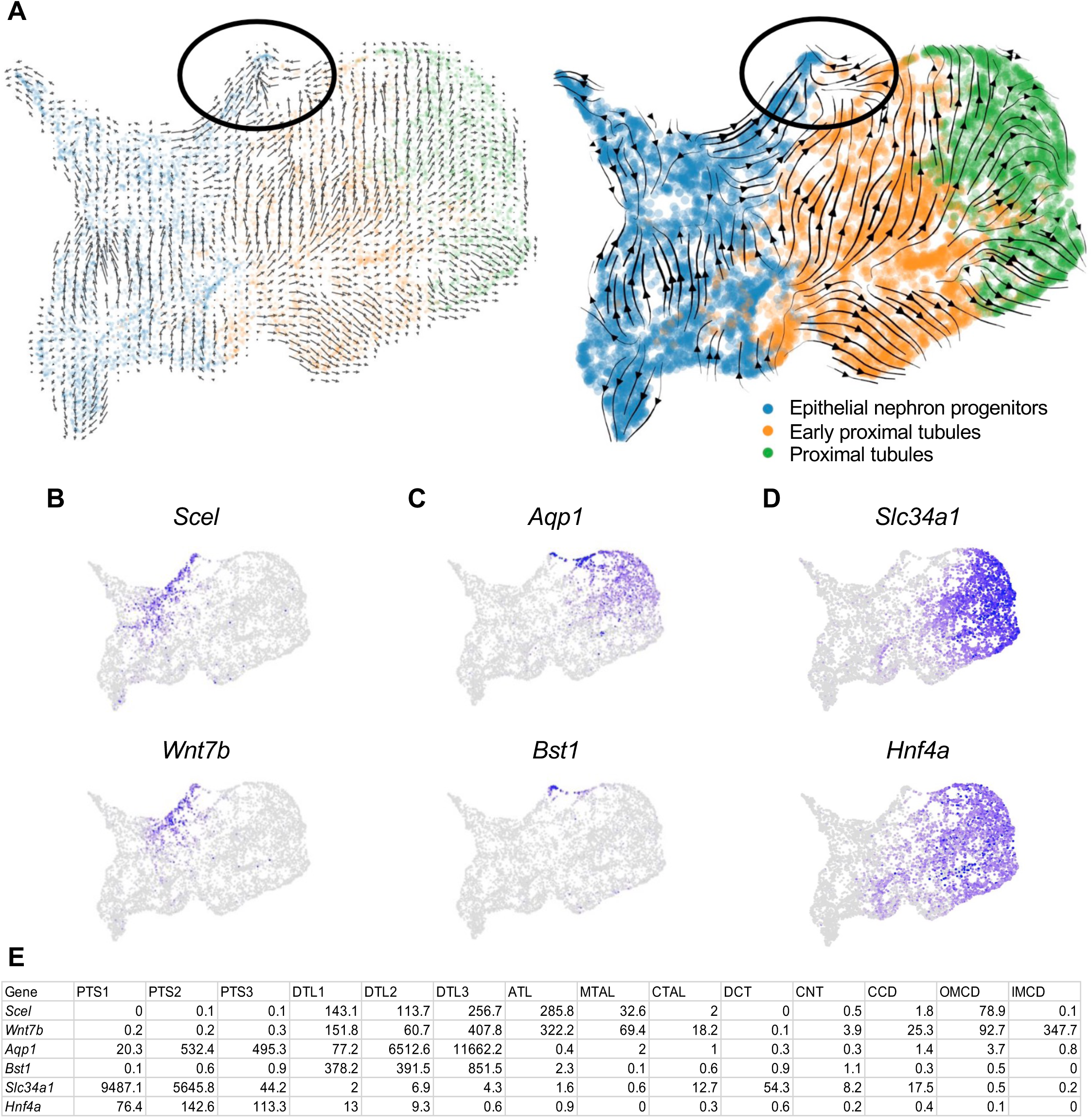
RNA velocity analysis. **(A)** Velocity grid plot and Velocity stream plot show that the descending limb (black ellipse) may emerge from Slc34a1+ Hnf4a+ proximal tubule cells and Wnt7b+ Scel+ epithelial nephron progenitor cells. **(B)** *Scel* and *Wnt7b* mark epithelial nephron progenitors that may develop into the descending limb. **(C)** *Aqp1* and *Bst1* mark the descending limb. **(D)** *Slc34a1* and *Hnf4a* mark proximal tubules. **(E)** The expression of the genes shown in B-D in mouse adult kidney (GSE150338). The numbers represent TPM (transcripts per million). In the nephron lineage, *Scel* and *Wnt7b* are highly expressed in thin descending and ascending limbs of the loop of Henle. *Aqp1* and *Bst1* are highly expressed in the thin descending limb of the loop of Henle. *Slc34a1* is highly expressed in the S1 and S2 segments of proximal tubules while Hnf4a is expressed in all three segments of proximal tubules. Abbreviations are listed in Supplemental Table 3.

**Supplemental Figure 8.**
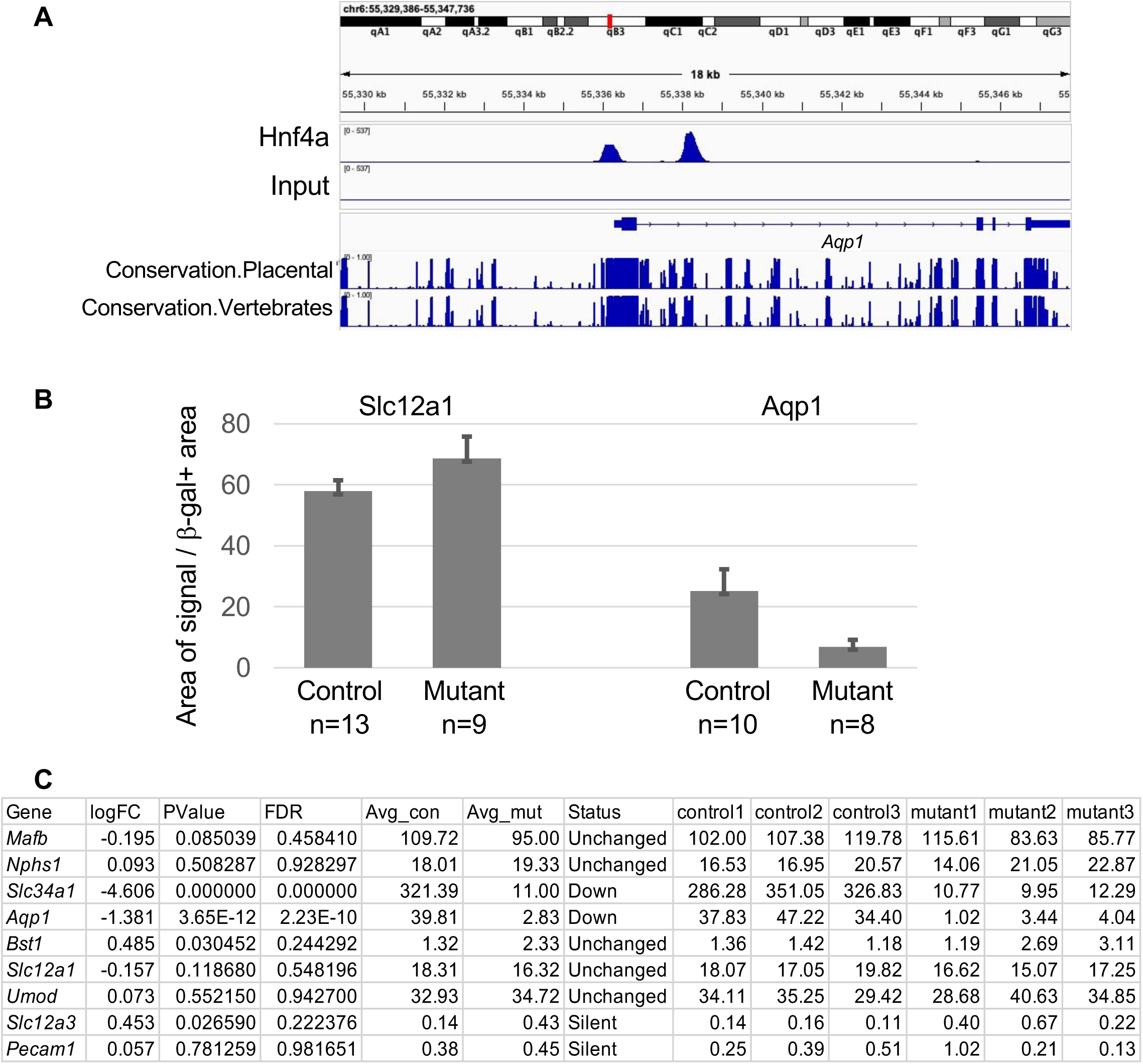
Regulation of *Aqp1* by Hnf4a. **(A)** Hnf4a directly binds to the promoter and a putative enhancer of *Aqp1*. Hnf4a chromatin immunoprecipitation sequencing was performed in newborn mouse kidneys (GSE144824). mouse genome, mm10. **(B)** Quantification of immunofluorescence images from Figure 4 shows that *Hnf4a* mutant kidneys exhibit an 18% increase in Slc12a1+ area (p = 0.0002, unpaired t-test) and a 73% reduction in Aqp1+ area (p = 0.0001, unpaired t-test) **(C)** Bulk RNA-seq analysis of FACS-isolated cells from the *Hnf4a* mutant and control kidneys at E18.5 (GSE291740). *Osr2Cre* was used to activate Rosa26 Ai3 reporter and delete Hnf4a. Because *Osr2Cre* does not target the distal tubule or endothelial cells, both *Slc12a3* (a distal tubule marker) and *Pecam1* (an endothelial marker) show minimal expression. Loss of Hnf4a leads to a significant downregulation of *Slc34a1* and *Aqp1*, suggesting that Hnf4a is essential for their expression. The expression of genes specific to podocytes (*Mafb* and *Nphs1*) or the thick ascending limb of the loop of Henle (*Slc12a1* and *Umod*) remains unaffected by the loss of Hnf4a. The full results are presented in Supplemental Table 4.

Supplemental Table 1. Primary antibodies used in this study

Supplemental Table 2. List of the marker genes for clusters identified in scRNA-seq dataset

Supplemental Table 3. Top 20 genes that are preferentially expressed in descending limb of loop of Henle in mouse adult kidney

Supplemental Table 4. RNA-seq analysis of FACS-isolated cells labeled by *Osr2Cre* from E18.5 *Hnf4a* control and mutant kidneys

## Supplemental Methods

### Study Design

For tissue immunofluorescence staining, we compared kidneys from control and mutant animals from the same litter. Control and mutant genotypes are labeled in each Figure. One experiment consists of kidneys from three embryos (two mutants, one control) or two animals (one mutant, one control) from one litter. We show representative results from at least three independent experiments from three separate litters, as indicated by (n=number) in Figure Legends. Sample size was determined by mating outcomes and size of litter. Kidneys from a litter were processed in parallel, in random order until genotype was determined. E.C. was aware of the group allocation (control or mutant genotype) from frozen block preparation stage to data analysis. Control and mutant samples were set in the same block so that a single section would contain both groups. A single section containing both control and mutant tissue underwent the same staining process as described in Methods and were imaged under the same exposure conditions in one sitting. Outcomes were defined by absence, presence, decrease or increase of signal and apparent differences in signal pattern.

The reference single cell RNA-seq dataset was constructed with 67,438 cells by combining seven separate scRNA-seq datasets of non-mutant mouse kidneys at embryonic day 18.5 (E18.5) or postnatal day 0 (P0). Batch correction was successful as the same cell types from each sample clustered together (Supplemental Figure 2).

### Antibodies and reagents used for immunostaining

Primary antibodies used in this study are listed in Supplemental Table 1. Secondary antibodies were Alexa Fluor® 647 Donkey Anti-Chicken IgY, (703-605-155), Alexa Fluor® 488-AffiniPure F(ab’)2 Fragment Donkey Anti-Chicken IgY (IgG) (703-546-155), Alexa Fluor® 647-AffiniPure F(ab’)2 Fragment Donkey Anti-Rabbit IgG (711-606-152), Alexa Fluor® 488 AffiniPure Donkey Anti-Guinea Pig IgG (706-545-148), Alexa Fluor® 647 AffiniPure Donkey Anti-Goat IgG (705-605-147), Alexa Fluor® 647 AffiniPure Donkey Anti-Rat IgG (712-605-153) purchased from Jackson ImmunoResearch Laboratories Inc. We also used secondaries Alexa Fluor® 555 donkey anti-rabbit IgG (A-31572) and Alexa Fluor® 647 Goat Anti-Mouse IgG1 (A-21240) from Invitrogen. We used sterile-filtered, heat-inactivated sheep serum (Equitech SS-0500HI) for blocking agent and Prolong Gold for mounting medium (Invitrogen P36930).

### Quantification in Supplemental Figure 8B

In Figure 4, we show cropped, representative immunofluorescence images of the loop of Henle from control and mutant kidneys. For quantification shown in Supplemental Figure 8B, we analyzed the full field of each group. We measured Slc12a1 and Aqp1 signals in 8-13 representative sectors of the medulla of the control and mutant kidneys using ImageJ software. By color thresholding, we selected and measured the area of β- galactosidase reporter+ Slc12a1+ (double positive) pixels (a) and the area of all β- galactosidase+ pixels (b). Relative expression of Slc12a1 was defined as (a)/(b)x100. Likewise, relative expression of Aqp1 was determined as the area of β-galactosidase+ Aqp1+ pixels divided by the area of all β-galactosidase+ pixels, multiplied by 100. The number of sectors measured for each group is shown as n=number in Supplemental Figure 8B.

