## Supplementary material for "Proximal tubule cells contribute to the thin descending limb of the loop of Henle during mouse kidney development": SuppTable3

### Supplemental Table 3

| Gene | PTS1 | PTS2 | PTS3 | DTL1 | DTL2 | DTL3 | ATL | MTAL | CTAL | DCT | CNT | CCD | OMCD | IMCD |
|---|---|---|---|---|---|---|---|---|---|---|---|---|---|---|
| *Fst* | 0.6 | 0 | 0 | 6.3 | 2814.6 | 3.2 | 2.2 | 0.1 | 0.1 | 0.1 | 0 | 0.1 | 1.3 | 0.3 |
| *Corin* | 0.1 | 0.4 | 0.9 | 926.5 | 0.6 | 0.7 | 1.2 | 0 | 0 | 0 | 0 | 0 | 0.5 | 0.2 |
| *Wnt11* | 0 | 0.1 | 0.1 | 383.1 | 6.2 | 0.6 | 0.6 | 0.1 | 0 | 0.1 | 0 | 0.1 | 0.4 | 0.1 |
| *Bst1* | 0.1 | 0.6 | 0.9 | 378.2 | 391.5 | 851.5 | 2.3 | 0.1 | 0.6 | 0.9 | 1.1 | 0.3 | 0.5 | 0 |
| *Angpt2* | 0.4 | 0.3 | 0.3 | 2348.5 | 482.1 | 611.3 | 6.5 | 1.8 | 0.9 | 0.3 | 0.4 | 0.8 | 5.7 | 5.4 |
| *Nr2e3* | 0.4 | 0 | 0 | 3 | 13.6 | 778.4 | 4.5 | 0.1 | 0.1 | 0 | 0.1 | 0.1 | 1.2 | 0.3 |
| *Igkc* | 0.3 | 0.2 | 0 | 6.6 | 5.4 | 217.4 | 0 | 0 | 0.1 | 1.1 | 0.5 | 0.3 | 0 | 0 |
| *Cntf* | 0 | 0 | 0.2 | 5.4 | 340 | 62.6 | 1.3 | 0.1 | 0.3 | 0.3 | 0.4 | 0.4 | 0.7 | 1.2 |
| *Uncx* | 0.3 | 0 | 0 | 162.9 | 0.8 | 1.6 | 0.6 | 0 | 0.7 | 0 | 0 | 0 | 1 | 0.3 |
| *Plk5* | 0.2 | 1 | 1.1 | 0.8 | 68.1 | 101.9 | 0.1 | 0.2 | 0 | 0.1 | 0.1 | 0.1 | 0.2 | 0.2 |
| *Pitx2* | 4.3 | 0.6 | 0.9 | 10.2 | 318.4 | 301.2 | 3.8 | 0.1 | 0.1 | 0.1 | 0.1 | 0.3 | 2.1 | 0.5 |
| *Cdh13* | 0.9 | 0.1 | 0.1 | 11.3 | 307.9 | 3.9 | 3 | 0.1 | 0.7 | 0.3 | 0.1 | 0.4 | 1.8 | 0.5 |
| *Dmkn* | 0.3 | 0 | 0 | 11.4 | 1.5 | 126.9 | 1.1 | 0 | 0 | 0 | 0 | 0 | 0.6 | 3.3 |
| *Sorcs3* | 0.1 | 0 | 0 | 32.4 | 2.2 | 69.3 | 4 | 0 | 0 | 0 | 0 | 0 | 0.1 | 0 |
| *Akap12* | 0.3 | 0.3 | 0.6 | 31.7 | 83.9 | 152.3 | 3.7 | 0.5 | 1.1 | 0.8 | 1.1 | 0.7 | 1.8 | 0.6 |
| *Tert* | 0.5 | 0.4 | 0.3 | 81.7 | 220.7 | 1343.5 | 26.7 | 0.5 | 0.5 | 0.5 | 1.3 | 1.8 | 5.2 | 35.3 |
| *Atoh8* | 0.3 | 0.2 | 0.1 | 93.8 | 10.5 | 0.6 | 1.9 | 0.9 | 0.3 | 0.3 | 0.3 | 0.1 | 0.7 | 0.3 |
| *Aqp1* | 20.3 | 532.4 | 495.3 | 77.2 | 6512.6 | 11662.2 | 0.4 | 2 | 1 | 0.3 | 0.3 | 1.4 | 3.7 | 0.8 |
| *Ifit3* | 4.2 | 2.1 | 3.6 | 2032.4 | 321 | 439.9 | 94 | 8 | 11.2 | 8.2 | 7.3 | 5.5 | 15.5 | 10.8 |
| *Cfap52* | 0.1 | 0.1 | 0.2 | 10.7 | 58 | 516.4 | 22.2 | 0.3 | 0.1 | 0.3 | 0.7 | 1.1 | 5.4 | 9 |

Top 20 genes that are preferentially expressed in descending limb of loop of Henle in mouse adult kidney (GSE150338). The numbers represent TPM (transcripts per million). After removing the genes whose TPM in DTL segments is less than 100, the rest of the genes were sorted based on the ratio of (TPM in DTL segments)/(TPM in all nephron segments).

PTS1, initial segment of the proximal convoluted tubule
PTS2, proximal straight tubule in cortical medullary rays
PTS3, last segment of the proximal straight tubule in outer stripe of outer medulla
DTL1, short descending limb of the loop of Henle
DTL2, long descending limb of the loop of Henle in the outer medulla
DTL3, long descending limb of the loop of Henle in the inner medulla
ATL, thin ascending limb of the loop of Henle
MTAL, medullary thick ascending limb of the loop of Henle
CTAL, cortical thick ascending limb of the loop of Henle
DCT, distal convoluted tubule
CNT, connecting tubule
CCD, cortical collecting duct
OMCD, outer medullary collecting duct
IMCD, inner medullary collecting duct
